# Human Cytomegalovirus UL138 Interaction with USP1 Activates STAT1 in infection

**DOI:** 10.1101/2023.02.07.527452

**Authors:** Kristen Zarrella, Pierce Longmire, Sebastian Zeltzer, Donna Collins-McMillen, Meaghan Hancock, Jason Buehler, Justin M. Reitsma, Scott S. Terhune, Jay A. Nelson, Felicia Goodrum

## Abstract

Innate immune responses are crucial for limiting virus infection. However, viruses often hijack our best defenses for viral objectives. Human Cytomegalovirus (HCMV) is a beta herpesvirus which establishes a life-long latent infection. Defining the virus-host interactions controlling latency and reactivation is vital to the control of viral disease risk posed by virus reactivation. We defined an interaction between UL138, a pro-latency HCMV gene, and the host deubiquintase complex, UAF1-USP1. UAF1 is a scaffold protein pivotal for the activity of ubiquitin specific peptidases (USP), including USP1. UAF1-USP1 sustains an innate immune response through the phosphorylation and activation of signal transducer and activator of transcription-1 (pSTAT1), as well as regulates the DNA damage response. After the onset of viral DNA synthesis, pSTAT1 levels are elevated and this depends upon UL138 and USP1. pSTAT1 localizes to viral centers of replication, binds to the viral genome, and influences UL138 expression. Inhibition of USP1 results in a failure to establish latency, marked by increased viral genome replication and production of viral progeny. Inhibition of Jak-STAT signaling also results in increased viral genome synthesis in hematopoietic cells, consistent with a role for USP1-mediated regulation of STAT1 signaling in the establishment of latency. These findings demonstrate the importance of the UL138-UAF1-USP1 virus-host interaction in regulating HCMV latency establishment through the control of innate immune signaling. It will be important going forward to distinguish roles of UAF1-USP1 in regulating pSTAT1 relative to its role in the DNA damage response in HCMV infection.

**Importance:** Human cytomegalovirus (HCMV) is one of nine herpesviruses that infect humans. Following a primary infection, HCMV establishes a life-long latent infection that is marked by sporadic, and likely frequent reactivation events. While these reactivation events are asymptomatic in the immune competent host, they pose important disease risks for the immune compromised, including solid organ or stem cell transplant recipients. Its complex interactions with host biology and deep coding capacity make it an excellent model for defining mechanisms important for viral latency and reactivation. Here we define an interaction with host proteins that commandeer typically antiviral innate immune signaling for the establishment of latency.

## Introduction

Viral latency is a remarkable *coup d’etat* that allows for the virus to persist in the face of robust antiviral immunity and is a hallmark of all herpesvirus infections. Latency is defined as a quiescent state of persistence whereby viral genomes are maintained in the absence of progeny virus production. During latency, viral gene expression is quieted, but not completely silenced [1-3]. The β-herpesvirus, human cytomegalovirus (HCMV), establishes latency in hematopoietic progenitor cells (HPCs within the CD34+ subpopulation) and cells of the myeloid lineage [4, 5]. HCMV latency is marked by sporadic and frequent asymptomatic reactivation events, which are controlled by the human immune response. In the absence of adequate cellular immunity, such as in the case of solid organ or hematopoietic cell transplantation, HCMV reactivation poses life threatening disease risk [5-7]. HCMV infection, reinfection or reactivation during pregnancy and subsequent transmission to the fetus makes HCMV the primary infectious cause of congenital birth anomalies [8, 9]. The benefits or consequences of lifelong persistence of HCMV in otherwise healthy individuals remain poorly defined [10-15]. Understanding the molecular basis of latency and reactivation is important in developing strategies to target or control the latent reservoir of HCMV and reveals fundamental mechanisms at the pinnacle of virus-host co-evolution.

HCMV encodes viral genes important for latency within the unique long *b*’ (UL*b*’) region of the HCMV genome. The UL*b*’ region is a contiguous 15-kilobase region spanning *UL132-UL150*. These genes are largely dispensable for replication in fibroblasts, but strongly impact infection outcomes in hematopoietic and endothelial cells [16, 17]. One of these genes, *UL138*, restricts virus replication in CD34+ HPCs and disruption of *UL138* results in a virus that replicates in HPCs in the absence of a stimulus for reactivation. UL138 functions, at least in part, in modulating the trafficking and signaling of cell surface receptors including, epidermal growth factor receptor (EGFR) [18, 19], tumor necrosis factor receptor-1 (TNFR1) [20, 21] and multidrug resistance-associated protein-1 (MRP1) [22]. UL138 sustains EGFR signaling, which drives the expression of transcription factors that affect viral gene expression from the *UL133-UL138* locus and latency [18], and sensitizes cells to TNF-α for reactivation [23, 24]. A gene co-regulated with UL138, UL135, functionally opposes UL138 in the regulation of EGFR trafficking, targeting it for degradation and attenuating signaling for reactivation [25, 26], but has not been shown to impact other UL138-regulated receptors. UL138 function suppresses immediate early gene expression and virus replication, giving it pro-latency properties [17, 27, 28].

In this study, we define an interaction between UL138 and USP1-associated factor 1 (UAF1) and ubiquitin specific peptidase 1 (USP1). UAF1 is also known as WD-repeat domain 48 (WDR48) protein. UAF1 is a scaffold or chaperone protein that is required for the activity of ubiquitin specific peptidases, USP1, USP12 and USP46 [29, 30]. USP1 stimulates the deubiquitylation and inactivation of proteins involved in the DNA damage response [31, 32].

USP1 also positively regulates innate pathways by enhancing and sustaining phosphorylation of signal transducer and activator of transcription 1 (pSTAT1) [33], an important activator of the antiviral type-1 interferon response. We hypothesized that UL138 may commandeer the UAF1-USP1 interaction to regulate pSTAT1 for latency. We find that sustained pSTAT1 during HCMV infection depends on both UL138 and USP1. Additionally, we show that pSTAT1 localizes to sites of viral DNA synthesis and transcription to regulate viral gene expression. Moreover, USP1 and pSTAT1 activity serve as repressors of viral replication for the establishment of latency in CD34+ HPCs. Collectively, our findings provide mechanistic insight into how HCMV has coopted the innate immune response through STAT1 signaling to regulate decisions to enter or exit latency.

## Results

### Defining UL138-UAF1 host interactions

We identified host interacting proteins for UL138 by immunoprecipitating UL138 fused in-frame with the Flag epitope tag from fibroblasts infected with the recombinant TB40/E strain, TB40/E-UL138_FLAG_, and defining proteins associated by mass spectrometry (IP-MS/MS). Interacting candidates that precipitated from a control pull down using the FLAG antibody and a lysate from cells infected with TB40/E lacking the FLAG epitope were subtracted from the interacting candidate data set. Top-ranking interacting candidates were based on peptide count and coverage, summarized in Table 1. IP-MS/MS peptides and data are provided for these candidates in Supplementary Information (SI) Table 1. This screen previously identified the lower-ranking interaction with EGFR [19]. UAF1/WDR48 (herein referred to as UAF1) was the top ranked UL138 interacting protein, followed by WDR20 and USP12. Two previous studies have identified were stained for USP1, pY701-STAT1, STAT1the UL138 interaction with UAF1 (referred to as WRD48), WDR20, and USP12 [34, 35], independently validating this work. We verified the UAF1 interaction with UL138 by overexpressing UAF1 fused in frame with a hemagglutinin (HA, UAF1_HA_) epitope tag and UL138 fused in frame with myc epitope tag (UL138_myc_). Immunoprecipitation (IP) of UAF1_HA_ co-precipitated UL138_myc_ (Fig. 1A).

**Table 1.**
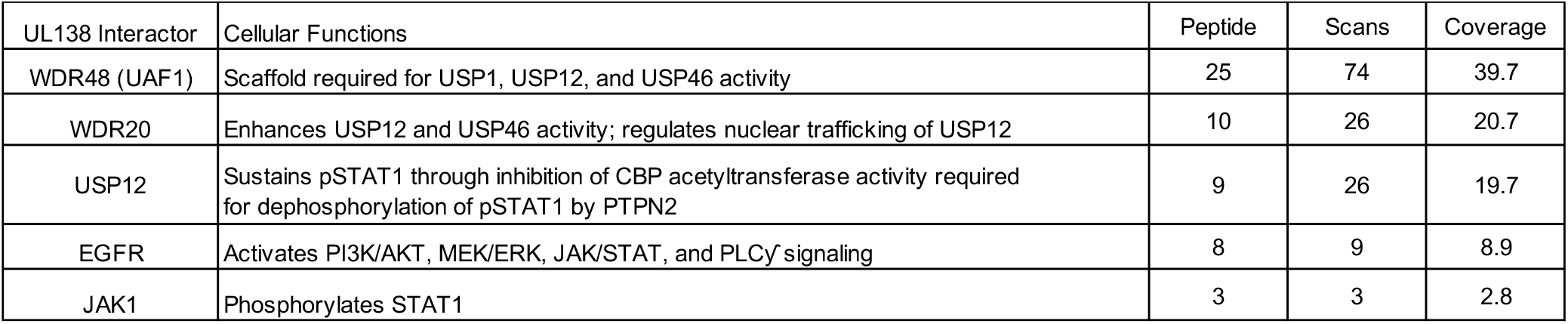
Summary of UL138 interactors. **HCMV latency protein UL138 interacts with UAF1 and associated proteins**. Fibroblasts were infected (MOI=1) with WT-*UL138*_FLAG_. At 48 hpi, *UL138*_FLAG_ was immunoprecipitated with a Flag antibody and following tryptic digest, peptides were identified by IP-MS/MS. Interacting candidates were subtracted from a control Flag pull-down from infected fibroblasts without a Flag tagged UL138. 128 candidates were identified and top interactions were identified by STRING and NCBI analysis.

**Fig 1.**
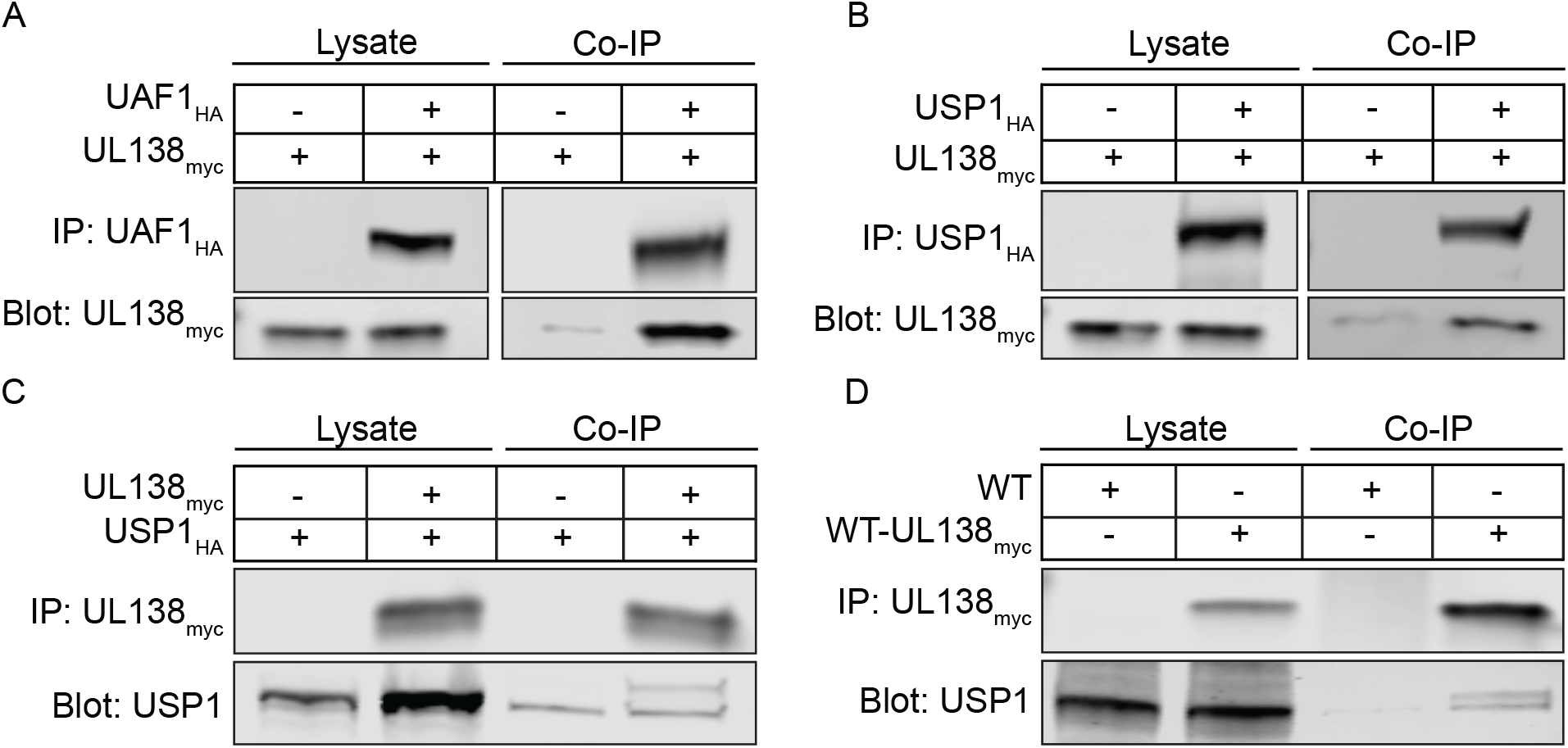
UL138 interacts with UAF1 and USP1. Human Embryonic Kidney (HEK293T) cells were transfected with a UL138_myc_ expressing plasmid along with either an empty vector control plasmid or with (A) UAF1_HA_ or (B) USP1_HA_. At 24 hours post transfection, the HA tag was immunoprecipitated. (C) A reciprocal immunoprecipitation where HEK293T cells were transfected with a USP1_HA_ expressing plasmid along with an empty vector control plasmid or a UL138myc expressing plasmid. USP1_HA_ was immunoprecipitated. (D) Fibroblasts were infected (MOI=1) with a WT-UL138_myc_ virus. At 24 hpi UL138_myc_ was immunoprecipitated.

USP1 is the most well defined UAF1 interactor, but likely to escape an interactome screen. To determine if USP1 is associated with UL138-UAF1 complexes, we exogenously expressed UL138_myc_ and USP1 fused in frame with a hemagglutinin (HA, USP1_HA_) epitope tag. We detected UL138-USP1 interaction through the co-immunoprecipitation of USP1_HA_ with UL138_myc_ (Fig. 1B), as well as the reciprocal IP (Fig. 1C). To determine if UL138 and USP1 interact during HCMV infection, cells were infected with a TB40/E-*UL138*_myc_ virus. At 24 hours post infection (hpi), UL138_myc_ IP co-precipitated USP1 (Fig. 1D). While not defined here, the interaction between UL138 and USP1 is likely indirect through UAF1. These results validate the interaction between UL138, UAF1, and USP1.

### UL138 sustains pY701-STAT1 during late times of infection

USP1 is well defined in its role in deubiquitinating monoubiquitylated PCNA [36] and Fanconi anemia pathways proteins, FANCD2 and FANCI [31, 32], to regulate the DNA damage response. Alternatively, a role for USP1 has been reported to enhance the activation of STAT1 (phosphorylation on tyrosine 701, pY701-STAT1) by deubiquitinating and stabilizing tank binding kinase 1 (TBK1) [33]. While the role of UL138 modulating the DDR will be addressed in a separate study, we hypothesized that UL138 may interact with and direct UAF1-USP1 to enhance and sustain pY701-STAT1 during HCMV infection in order to restrict virus replication.

To explore this, we analyzed pY701-STAT1 in fibroblasts infected with wild type TB40/E (WT) or a virus containing stop codon substitutions in its 5’ end to disrupt UL138 protein synthesis, TB40/E-*UL138*_STOP_ (Δ*UL138*_STOP_) [37]. Over a time course of 72 hpi, we observed enhanced and sustained pY701-STAT1 in the WT infection compared to the Δ*UL138*_STOP_ infection (Fig. 2A). STAT1 levels increased and were sustained with WT infection. By contrast, pY701-STAT1 was lost in the Δ*UL138*_STOP_ infection and this was accompanied by a diminishment in STAT1 levels to resemble uninfected cells by 48 hpi. The diminishment of total STAT1 may be the result of feedback in STAT1 signaling. STAT1 transcripts and protein levels are increased by pY701-STAT1 signaling [38] and could account for the increase in STAT1 levels detected in the WT infection. Multiple independent experiments are quantified in Figure 2B. Immediate early (IE) proteins accumulated to greater levels in Δ*UL138*_STOP_ infection relative to the WT infection, consistent with increased IE gene expression and replication of a UL138-mutant virus infection [17, 39]. These results suggest UL138 enhances and sustains pY701-STAT1 levels during WT infection.

**Fig 2.**
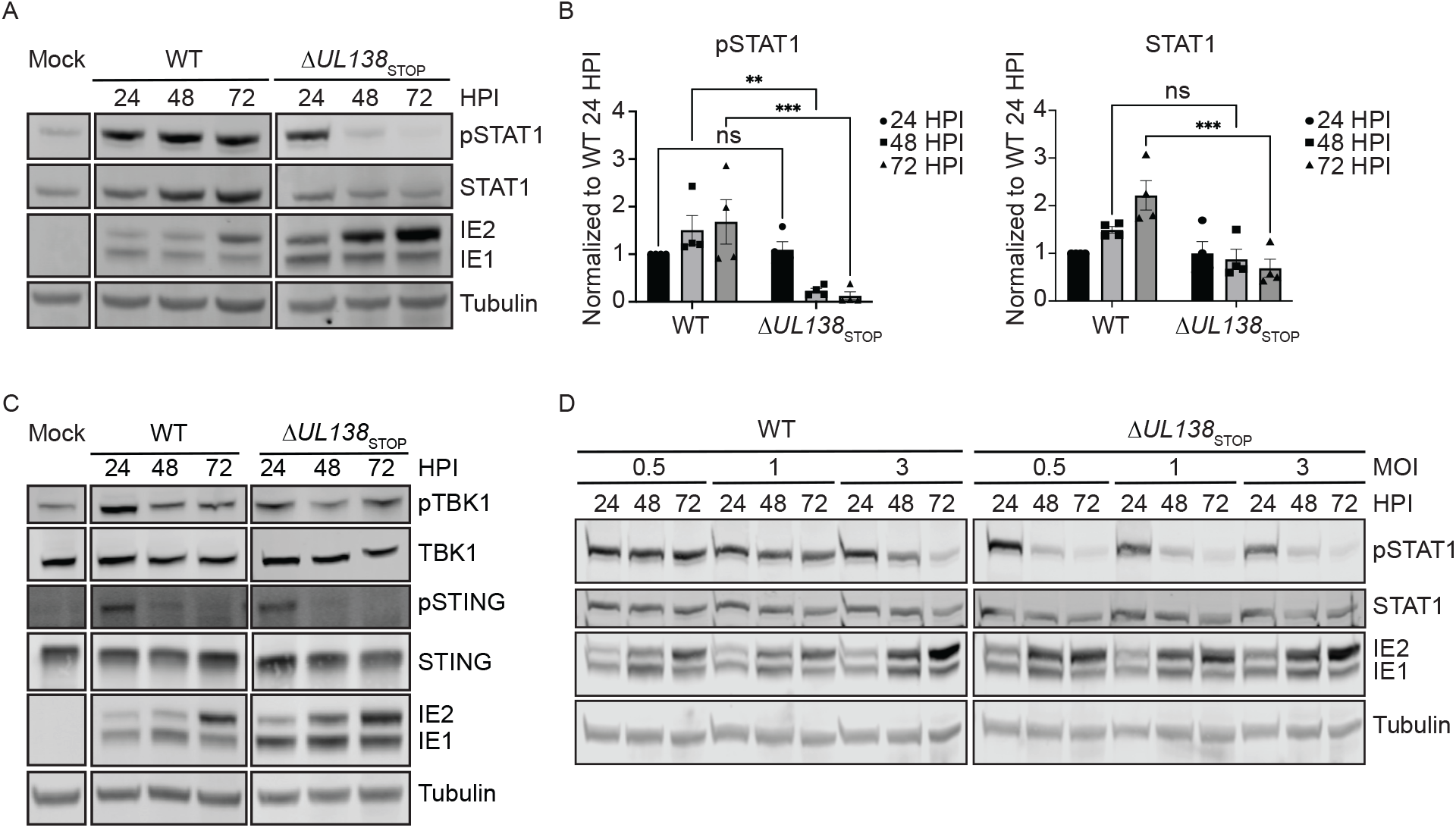
UL138 enhances and sustains pY701-STAT1 during late times of infection. (A) Fibroblasts were infected (MOI=1) with a WT or Δ*UL138*_STOP_ virus for 24-72 hpi. At each time point, lysates were lysed and immunoblotted. Blots were stained with pY701-STAT1, STAT1, IE1/2 antibody, and Tubulin. IE proteins serve as a control for infection and tubulin serves as a control for loading. (B) pY701-STAT1 and STAT1 were normalized to WT 24 hpi and graphed to calculate statistics. Statistical significance was calculated by Two-Way Anova and represented by asterisks. Graphs represent the mean of three replicates and error bars represent SEM. (C) Fibroblasts were infected (MOI=1) with either WT or Δ*UL138*_STOP_ virus and lysates were immunoblotted. Blots were stained with pTBK1, TBK1, pSTING, STING, IE1/2, and Tubulin. (D) Fibroblasts were infected at MOI of 0.5, 1, and 3 with a WT or Δ*UL138*_STOP_ virus and lysates were immunoblotted. Blots were stained for pSTAT1, STAT1, IE1/2, and Tubulin.

Given the reported roles of USP1 in stabilizing TBK1 [40] and of UL138 in targeting STING for degradation outside the context of infection [41], we analyzed total and phosphorylated levels of TBK1 and STING during HCMV infection. Phosphorylated and total levels of TBK1 and STING remained unchanged between WT and Δ*UL138*_STOP_infections (Fig. 2C). Additionally, TBK1 levels remained unchanged with chemical inhibition of USP1 with C527 (SI Fig. 1). From these results, we conclude that UL138 regulates a late phase induction of pY701-STAT1 during HCMV infection independently of TBK1 and STING regulation.

STAT1 is activated by type-1 interferon paracrine signaling. Therefore, it is possible that the enhanced pY701-STAT1 detected in WT infection is indirectly due to UL138 regulation of type-I interferon secretion and activation of STAT1 in neighboring uninfected cells. To investigate this, we infected fibroblasts with a WT or Δ*UL138*_STOP_ virus at increasing multiplicities of infection (MOI) (Fig. 2D). At an MOI of 0.5 or 1, the WT infection augmented pY701-STAT1 across the 72 hpi time course, although the activation of pY701-STAT1 was overcome at an MOI of 3 by 72 hpi. Strikingly, the Δ*UL138*_STOP_ infection showed a stark loss of pY701-STAT1 regardless of MOI, suggesting UL138 is required for late phase induction of pY701-STAT1 in a paracrine-independent manner. The loss of pY701-STAT1 in the WT infection at high MOI may reflect the action of multiple viral negative regulators on innate signaling [42] that override UL138-mediated induction of pSTAT1 during a replicative infection, particularly at high multiplicities.

### Late phase induction of pY701-STAT1 depends on USP1

To determine if the UAF1-USP1 interaction was involved in UL138-mediated regulation of pY701-STAT1, we inhibited UAF1-USP1 interactions with the chemical inhibitor, C527 [43] in fibroblasts infected with WT or Δ*UL138*_STOP_ viruses. Relative to DMSO, C527 treatment dramatically diminished pY701-STAT1 levels in cells infected with WT virus but had little effect on Δ*UL138*_STOP_ infection (Fig. 3A), suggesting a role for USP1 in the UL138-associated induction of pY701-STAT1. We further confirmed the role of USP1 in the UL138-mediated induction of pY701-STAT1 through siRNA-mediated USP1 knockdown (Fig. 3B). Relative to a non-targeting control (NTC) specific to luciferase, we achieved approximately 80% knockdown of USP1 (Fig. 3C). Consistent with USP1 inhibition, USP1 knockdown diminished pY701-STAT1 in WT infection but had little to no effect on pY701-STAT1 in Δ*UL138*_STOP_ infection (Fig. 3D).

**Fig 3.**
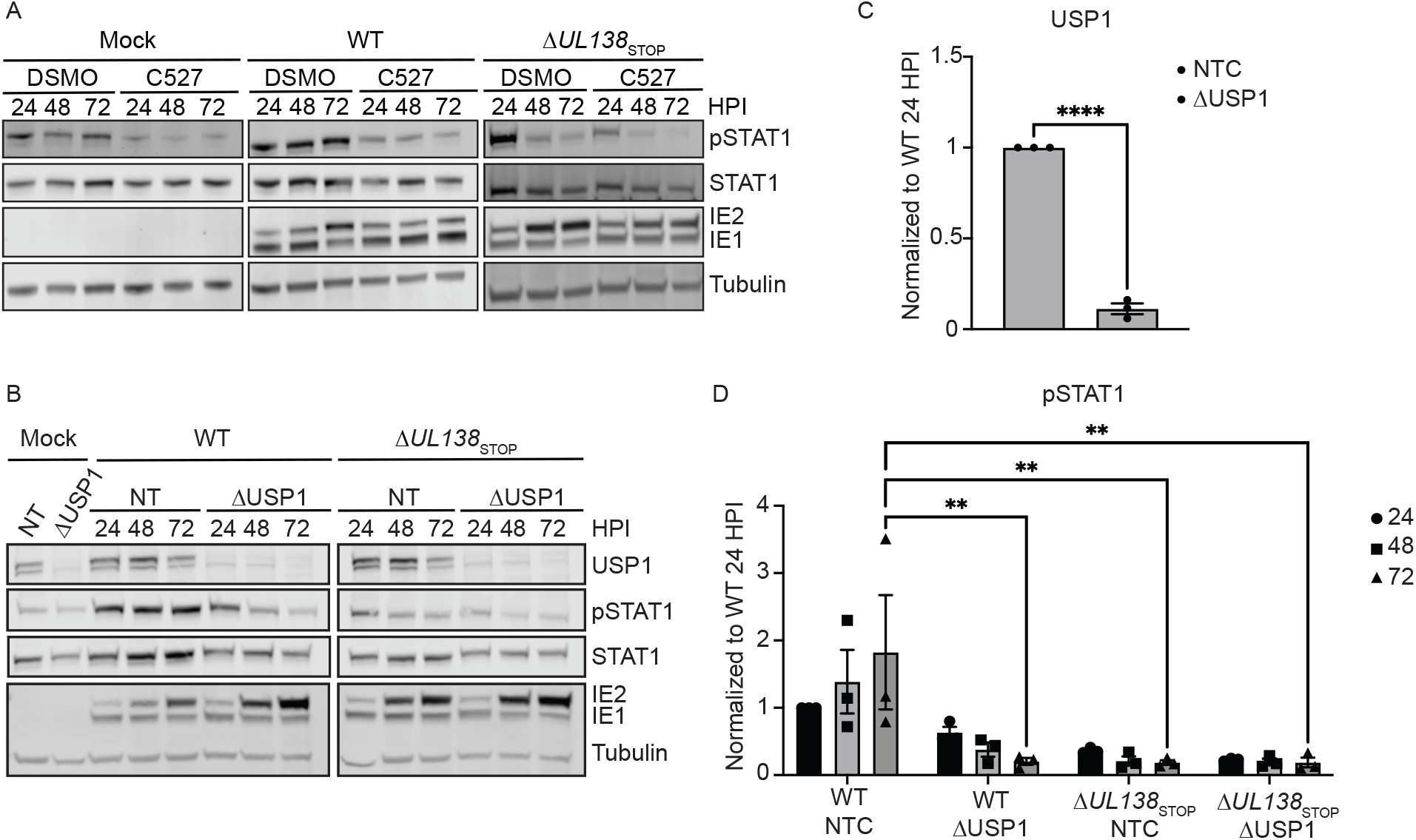
UL138 induction of pY701-STAT1 depends on USP1 activity. Fibroblasts were treated with either DMSO (vehicle control) or C527 24 hours prior to being infected (MOI=1) with a WT or Δ*UL138*_STOP_ virus. Lysates were collected and each timepoint and immunoblotted. Blots were stained with pY701-STAT1, STAT1, IE1/2, and Tubulin. (B) Fibroblasts were reverse transfected with 3 combined siRNA for a non-targeting control (NTC) or USP1. 24 hours post reverse transfection, the media was changed. 48 hours post reverse transfection, fibroblasts were infected (MOI=1) with either a WT or Δ*UL138*_STOP_ virus and lysates were immunoblotted. Blots were stained for USP1, pY701-STAT1, STAT1, IE1/2, and Tubulin. (C). USP1 levels were normalized to NTC and graphed to calculate statistics. Statistical significance for USP1 was calculated using an unpaired t test and represented by asterisks. (D) pY701-STAT1 was normalized to WT NTC 24 hpi and graphed to calculate statistics. Statistical significance was calculated by Two-Way Anova and represented by asterisks. Graphs represent the mean of three replicates and error bars represent SEM.

### UL138 enhances an early ISG response

Phosphorylation and activation of STAT1 drives its translocation to the nucleus to regulate the expression of hundreds of ISGs [38]. Therefore, we next analyzed pY701-STAT1 localization by subcellular fractionation experiments. pY701-STAT1 accumulated in the nuclei of cells infected with either WT or Δ*UL138*_STOP_ infection at 24 hpi (Fig. 4A). However, a dramatic loss of pY701-STAT1 from the nuclei of Δ*UL138*_STOP_-infected cells was observed at 72 hpi. Taken together, UL138 enhances and sustains pY701-STAT1 and its localization to the nucleus at late times in infected fibroblasts.

**Fig 4.**
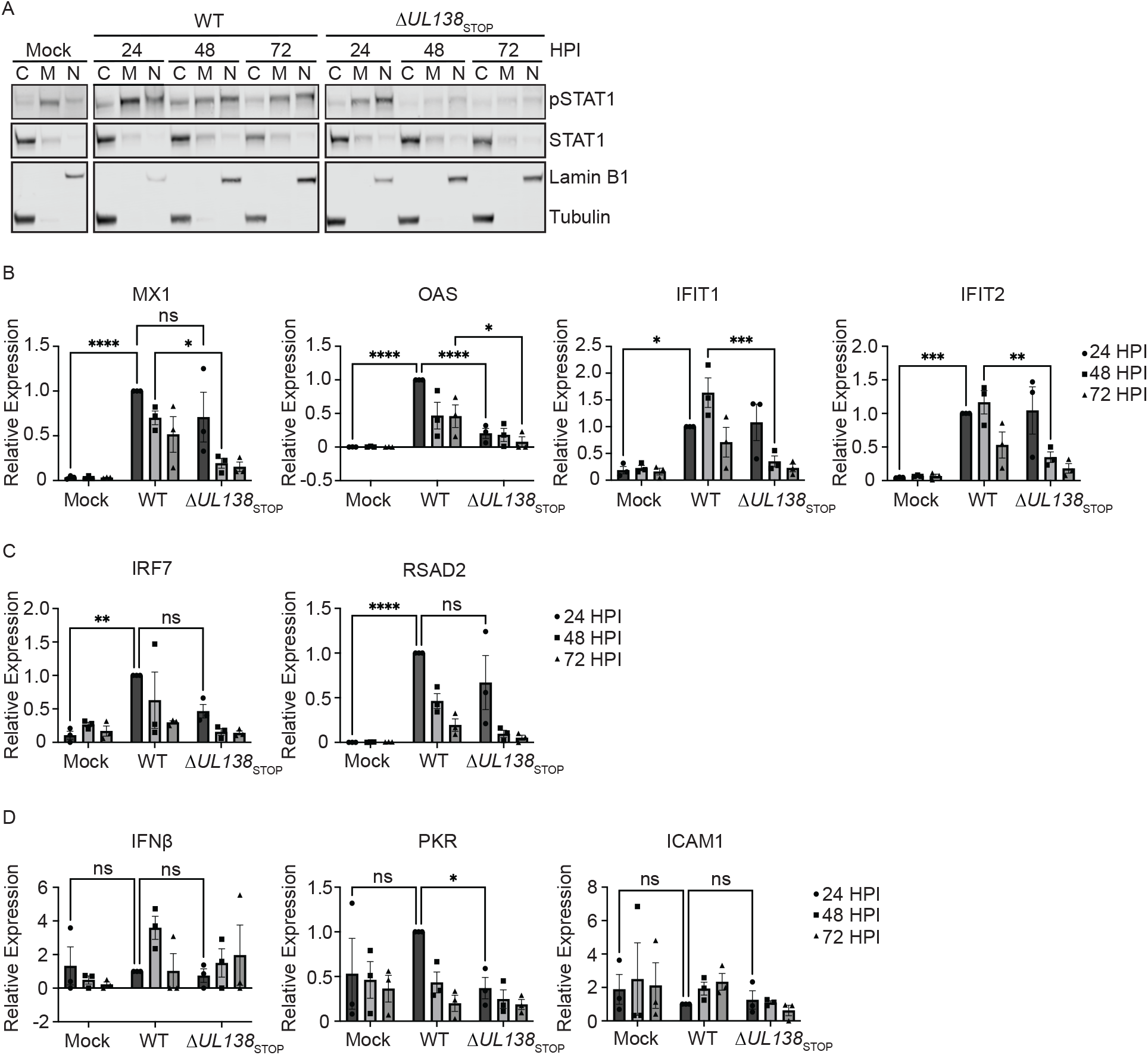
UL138 upregulates an early ISG response. (A) (B, C, D) Fibroblasts were infected (MOI=1) with a WT or Δ*UL138*_STOP_ virus. DNA was isolated at 24, 48, and 72 hpi. RT-qPCR for ISG expression was performed with TaqMan Gene Expression Assays (Thermo Fischer Scientific). Relative expression was determined using the ΔCT method and values were normalized to the WT 24 hpi and graphed for statistical analysis. Statistical significance was calculated using Two-Way Anova and represented by asterisks. Graphs represent the mean of three replicates and error bars represent SEM.

We next analyzed the induction of multiple interferon stimulated genes (ISGs) directly downstream of pY701-STAT1 in WT and Δ*UL138*_STOP_ infection by quantitative reverse transcriptase PCR (RT-qPCR). First, we analyzed ISGs induced by a type-1 IFN response through the formation of STAT1-STAT2 heterodimers. MX1, OAS, IFIT1, and IFIT2 showed a significantly larger induction in the WT-infection compared to the Δ*UL138*_STOP_-infection either at 24 hpi for OAS or 48 hpi for MX1, IFIT1 and IFIT2 (Fig. 4B). IRF7 and RSAD2 were similarly induced in both infections at 24 hpi compared to uninfected cells (Fig. 4C). Whereas IFNβand PKR were not induced with either infection above uninfected cells (Fig. 4D). ICAM1contains only a gamma activated sequence (GAS) for the binding of STAT1 homodimers when activated by the type-2 IFN pathway [44]. The expression of ICAM1 showed no induction compared to uninfected cells, consistent with a predominantly type-1 interferon response. These results suggest that UL138 sustains a specific type-1 IFN-response in HCMV infection, especially during the early times of infection.

### pY701-STAT1 localizes to sites of viral DNA synthesis and transcription

Localization of pY701-STAT1 to the nucleus during WT infection stimulates production of multiple ISGs early in infection. However, by 72 hpi, many of the induced ISGs return to similar levels seen in uninfected cells independently of the enhanced and sustained pY701-STAT1 seen at 72 hpi.Therefore, we questioned whether pY701-STAT1 was localizing to sites of viral DNA replication and transcription at late times of infection. We performed immunofluorescence for pY701-STAT1 on fibroblasts infected with a WT or Δ*UL138*_STOP_ virus at 72 hpi (Fig. 5A). Uninfected fibroblasts were untreated or treated with 1000U/mL of universal IFN for 30 minutes to induce pY701-STAT1 localization to the nucleus, as a positive control. In WT infected fibroblasts, pY701-STAT1 localized within the nuclei of infected cells and co-localized predominantly with UL44, a viral processivity factor that marks replication compartments (RCs), sites of viral DNA replication and transcription. There was no detection of pY701-STAT1 in a Δ*UL138*_STOP_ infection, consistent with reduced pY701-STAT1 protein levels (Fig. 2A). Uninfected neighboring cells in the WT infection showed a lack of induction of pY701-STAT1 and its localization to the nucleus, consistent with the conclusions that elevated pY701-STAT1 in a WT infection is not due to paracrine IFN signaling. Colocalization between UL44 and pY701-STAT1 was detected as early as 48 hpi in WT infection (SI Fig. 4). From these results, we conclude that UL138 sustains pY701-STAT1 and that pY701 predominantly localizes to viral RCs. This suggests a role for pSTAT1 in regulating viral gene expression, perhaps to the detriment of cellular ISG expression.

**Fig 5.**
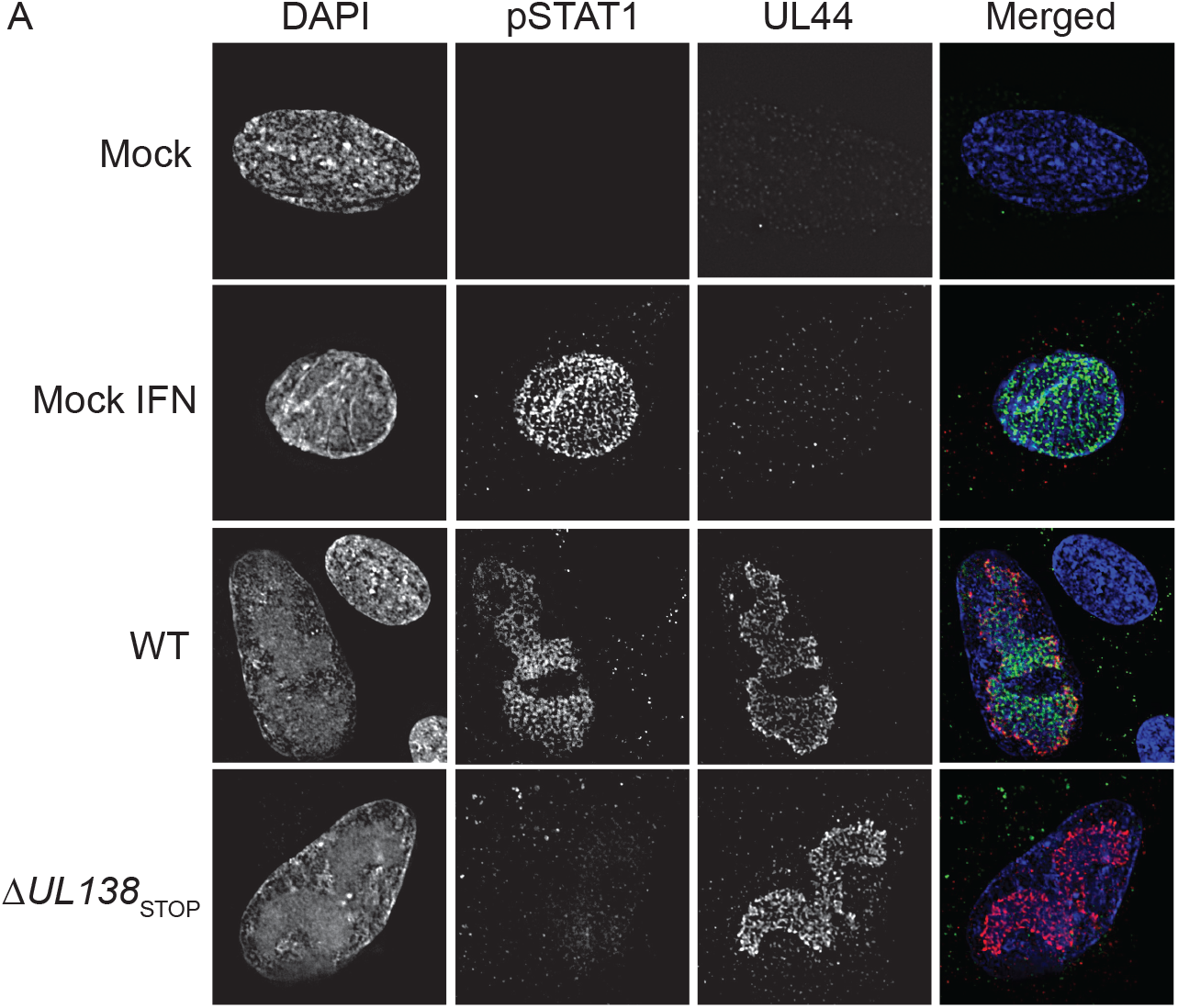
pY701-STAT1 localizes to sites of viral DNA synthesis and transcription at late times post infection. (A) Fibroblasts were plated on coverslips at 10,000 cells per coverslip. Fibroblasts were infected (MOI=1) with either a WT or Δ*UL138*_STOP_ virus for 72 hpi. At 72 hpi, cells were processed per the antibody manufacturer’s recommendations. Coverslips were imaged using a DeltaVision deconvolution microscope.

### Late phase pY701-STAT1 binds to the viral genome and regulates UL138 transcription

Considering the presence of pY701-STAT1 at RCs at 72 hpi, we were interested in whether pY701-STAT1 is binding to the viral genome as a transcription factor. Utilizing PhysBinder, we identified 86 pY701-STAT1 consensus sites on the viral genome. Two of which were within the UL133-UL138 locus upstream of UL138, designated site 1 and site 2 (Fig. 6A). To understand if pY701-STAT1 was binding to either of these two sites during HCMV infection, we performed CUT&RUN paired with RT-qPCR to quantitatively detect target sequences bound by STAT1 (Fig. 6B). CUT&RUN captures DNA bound by a protein using an antibody specific to the protein and micrococcal nuclease conjugated to protein A-protein G to cleave and release the DNA-protein complex [45, 46]. After interferon stimulation, pY701-STAT1 binds to an interferon-sensitive response element (ISRE) consensus sequence upstream of many ISGs to induce gene expression, including IRF1. We analyzed pY701-STAT1 binding to the IRF1 consensus sequence in uninfected fibroblasts with or without universal interferon stimulation for 30 minutes. In unstimulated cells, very low levels of pY701-STAT1 were bound to the IRF1 consensus site. However, upon universal interferon stimulation, there was a significant increase in pY701-STAT1 binding to the IRF1 consensus sequence. In WT infection at 48 hpi, pY701-STAT1 bound both site 1 and 2 upstream of UL138 relative to the uninfected IRF1 control, whereas we did not detect binding of STAT1 to these sites in Δ*UL138*_STOP_ infection. Therefore, this data suggests pY701-STAT1 binds to the UL133-UL138 region of the viral genome during HCMV infection in a manner dependent on UL138.

**Fig 6.**
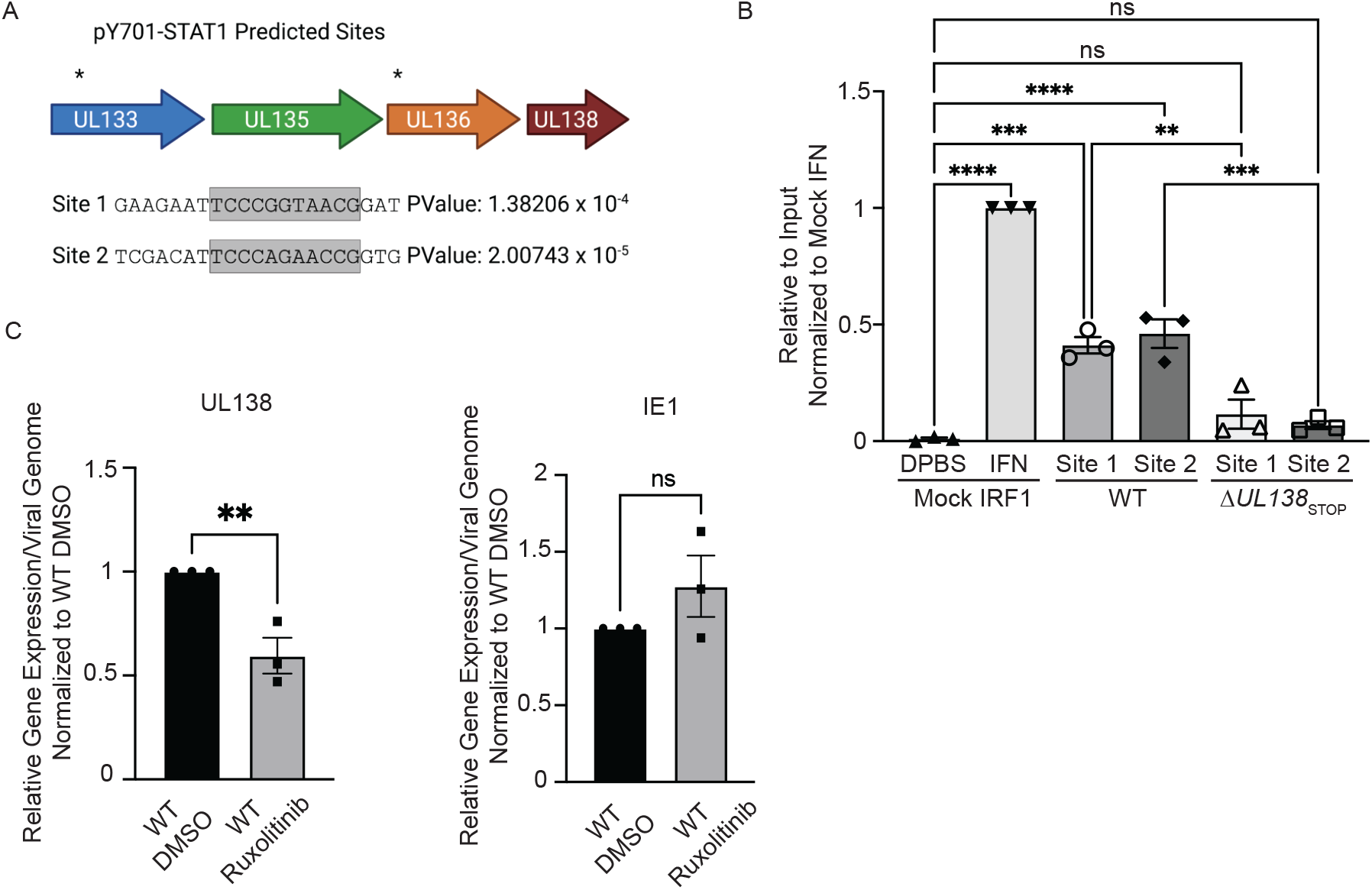
pY701-STAT1 binds to the viral genome and regulates UL138 transcription during a replicative infection. (A) The HCMV genome was run through PhysBinder Software to obtain potential pY701-STAT1 consensus sites on the viral genome. Site 1 and 2 are within UL133 and UL136 respectively and are upstream of UL138. (B) Fibroblasts were infected (MOI=1) with either a WT or Δ*UL138*_STOP_ virus. At 72 hpi, uninfected cells were treated with DPBS (vehicle control) or 1000U/ml of universal interferon for 30 minutes. After 30 minutes, all cells entered the CUT&RUN protocol developed by Cell Signaling Technologies. qPCR was performed on the input DNA and immunoprecipitated DNA utilizing primers specific to pY701-STAT1 binding sites of interest. Percent Immunoprecipitated was calculated to 2% of the input sample and graphed normalized to mock cells with interferon treatment. Statistical significance was calculated using One-Way Anova and represented by asterisks. (C) Fibroblasts were infected (MOI=1) with either a WT or Δ*UL138*_STOP_ virus. At the time of infection, cells were treated with DMSO (vehicle control) or 5uM Ruxolitinib every 24 hours. At 72 hpi, DNA and RNA was isolated and transcripts encoding UL138 and IE were quantified by RT-qPCR normalized to H6PD. Viral genomes were analyzed through qPCR using primers for b2.7kb and RNaseP as a loading control. Relative expression was determined using the ΔCT method and values were normalized to the DMSO control and graphed for statistical analysis. Statistical significance was calculated using unpaired student t test and represented by asterisks. Graphs represent the mean of three replicates and error bars represent SEM.

Given pSTAT1 binds to the viral genome at two sites upstream of UL138 during HCMV infection, we analyzed UL138 transcripts in infected fibroblasts in the presence of Ruxolitinib, a JAK1/2 inhibitor. Through RT-qPCR, we analyzed UL138 expression relative to viral genomes when phosphorylation of STAT1 was inhibited (Fig. 6C). With inhibition of JAK1/2, we found a 25% diminishment in UL138 gene expression but no effect on immediate early 1 (IE1) expression per viral genome in a TB40/E infection relative to a vehicle control. This data suggests that pY701-STAT1 binding to the viral genome regulates UL138 gene expression.

### USP1 and pY701-STAT1 are required for the establishment of viral latency

Given UL138 has been defined as important to the establishment of latency, we next wanted to investigate the possible role of USP1 in latency establishment. We chemically inhibited USP1 activity with C527 in primary CD34+ HPCs infected with WT or Δ*UL138*_STOP_. For these experiments, infected CD34+ HPCs were isolated by fluorescent activated cell sorting and then seeded into long term bone marrow cultures over a stromal cell support to maintain HPC phenotype and function [47]. At 10 dpi, HPC cultures were split. Half the cells were seeded by limiting dilution onto fibroblast monolayers in a cytokine-rich media to promote myeloid differentiation and HCMV reactivation. The other half of the cells were mechanically lysed and seeded in parallel by limiting dilution to determine virus produced during the latency culture prior to stimulation of reactivation (pre-reactivation). Fourteen days post seeding, infectious centers (GFP+ wells) were counted to determine the frequency of reactivation using extreme limiting dilution assay (ELDA). The WT virus establishes latency indicated by the low level of infectious centers measured in the pre-reactivation control and the frequency of reactivation increased following stimulation of reactivation. C527 inhibition of USP1 during latency culture increased virus replication and production of infectious centers in the pre-reactivation control to levels equivalent to the reactivation and the vehicle (DMSO) control-treated cells in three independent experiments, indicating a failure to establish latency (Fig. 7A). Δ*UL138*_STOP_ infection, as shown previously, also fails to establish a latent infection and replicates similarly to the WT infection with C527 treatment. Addition of C527 to Δ*UL138*_STOP_ infection had no additional effect on the phenotype, consistent with the UL138-USP1 interaction being a key function of UL138 in the establishment of latency. Three independent replicates are shown, but the experiments did not reach statistical significance. The Δ*UL138*_STOP_ infections show great variability depending on donor and therefore, the values were log2 transformed for graphing. This stabilizes the large variability associated with Δ*UL138*_STOP_ infections and allows us to better interpret the differences between WT DMSO pre-reactivation and the other conditions.

**Fig 7.**
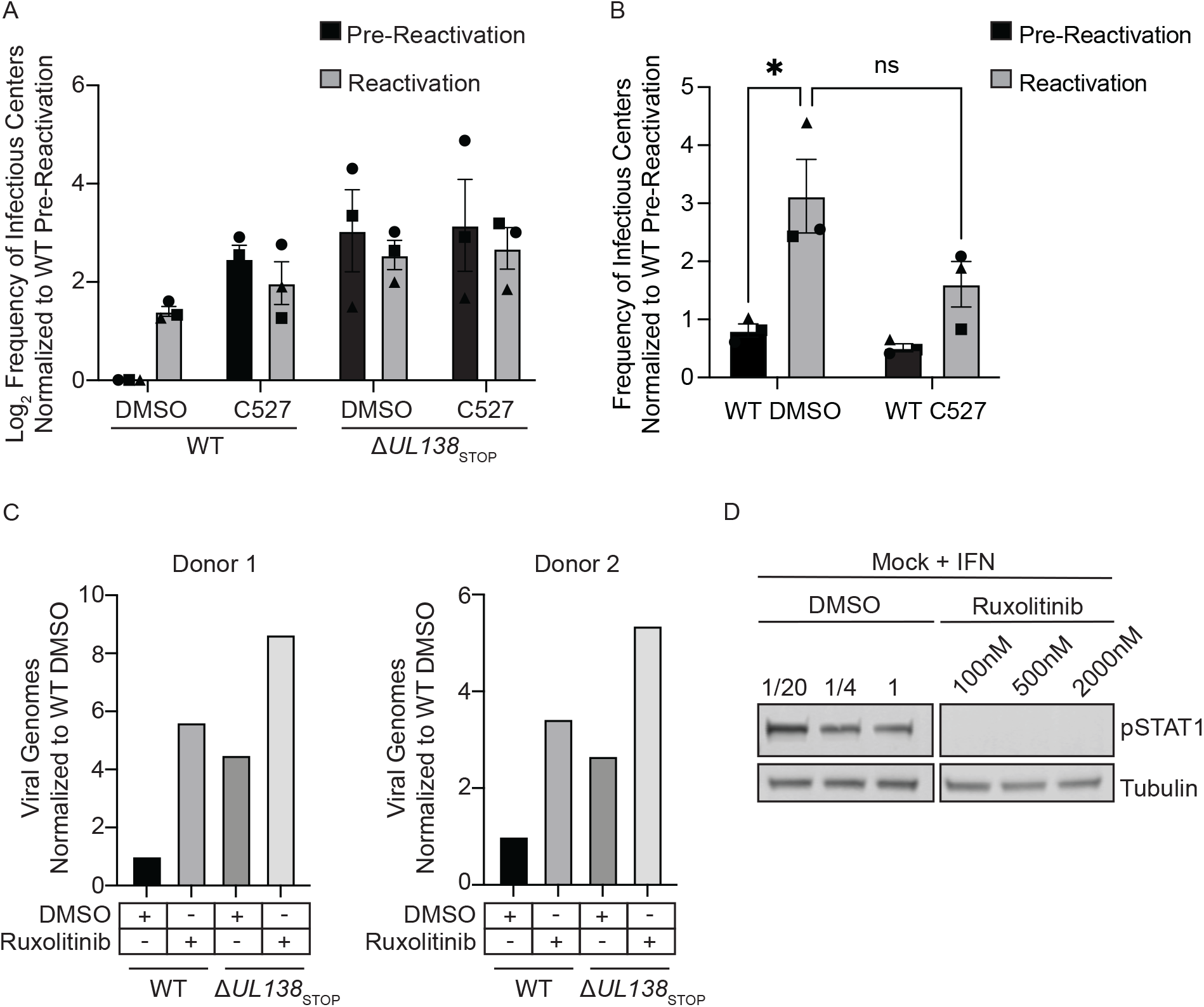
USP1 and pY701-STAT1 are required for suppression of viral replication and latency establishment. (A) CD34+ HPCs were infected with either WT or Δ*UL138*_STOP_ virus (MOI=2). At 24 hpi, CD34+/GFP+ cells were sorted and seeded into long-term culture in the presence of DMSO or 1uM C527. After 10 days in culture, populations were either mechanically lysed (pre-reactivation) or whole cells (reactivation) were plated onto fibroblast monolayers in cytokine-rich media. 14 days later, GFP+ wells were scored and frequency of infectious centers was determined by limited dilution analysis. The frequency of infectious centers was normalized to WT pre-reactivation and the average of 3 independent experiments is shown. Due to donor variability in the Δ*UL138*_STOP_ infection, data was Log2 transformed and graphed. (B) CD34+ HPCs were infected and sorted for long term culture as described in A without the addition of C527. At day 10, CD34+ HPCs were plated onto fibroblast monolayers in cytokine-rich media in the presence of DMSO or 1uM C527. 14 days later, GFP+ wells were scored and frequency of infectious centers was determined by limited dilution analysis. The frequency of infectious centers was normalized to WT pre-reactivation and the average of 3 independent experiments is shown. Statistical significance was calculated using Two-Way Anova and represented by asterisks. Graphs represent the mean of three replicates and error bars represent SEM. (C) CD34+ HPCs were infected with either WT or Δ*UL138*_STOP_ virus (MOI=2) in cytokine-rich media in the presence of DMSO or 500nM Ruxolitinib. Every 24 hours, 500nM Ruxolitinib was refreshed in the media. At days 1 and 5 post infection, DNA was isolated for qPCR using primers for b2.7kb and RNaseP. Viral genomes were determined using ΔCT method and the day 5 values were normalized to the corresponding day 1 values. For graphing, day 5 genomes were normalized to WT DMSO. Graphs represent cells from bone marrow donor 1 and donor 2 and error bars represent SEM. (D) To ensure Ruxolitinib inhibited pY701-STAT1, CD34+ HPCs were cultured in the presence of 100nm, 500nM, or 2000nM Ruxolitinib or equivalent DMSO for 3 hours before being treated with 1000U/ml universal interferon for 30 minutes and immunoblotted for pSTAT1 and tubulin.

As USP1 was important for the induction of STAT1 in fibroblasts (Fig. 3 A, B), we were interested in a possible role for USP1 specifically in viral reactivation. We infected CD34+ HPCs with a WT virus, as described above; however, we added C527 or DMSO as a vehicle control at the time of reactivation. Inhibition of USP1 at the time reactivation did not impact infectious centers production (Fig. 7B). Therefore, USP1 activity is important for latency establishment but not involved in reactivation.

Given the role of USP1 activity in the establishment of latency and the role of the type-I IFN response in reversible silencing of the genome [48], we next wanted to investigate the role of STAT1 in the establishment of latency. We infected CD34+ HPCs with a WT or Δ*UL138*_STOP_ virus in the presence or absence of the JAK1/2 inhibitor, Ruxolitinib, added at the time of infection. As reported previously [26], the failure of Δ*UL138*_STOP_ to enter latency is reflected in increased viral genome synthesis in CD34+ HPCs relative to the WT infection. Inhibition of JAK1/2 resulted in increased in viral genomes in the WT infection at 5 dpi, reflecting increased viral activity to the level of Δ*UL138*_STOP_ infection, indicating a failure to establish latency (Fig.7C). Ruxolitinib also increased genome replication in Δ*UL138*_STOP_ infection, but more modestly than in the WT infection. Experiments from two independent CD34+ HPC donors are shown. To ensure Ruxolitinib was inhibiting the phosphorylation of STAT1, uninfected CD34+ HPCs were treated with universal interferon in the presence of differing doses of DMSO or Ruxolitinib (Fig. 7D). JAK1/2 inhibition diminished pY701-STAT1 at all doses, however, 500nM was chosen because it was the highest dose that didn’t alter proliferation and survival of CD34+ HPCs. Taken together, these data suggest that USP1 and pY701-STAT1, as modulated by UL138, are important for repression of viral replication and successful establishment of viral latency.

## Discussion

Type 1 interferons include IFNαand IFNβ, which bind to IFNAR and stimulates JAK1/2-mediated phosphorylation of STAT1 and STAT2 along with recruitment of IRF9 to generate the ISGF3 complex [49]. The ISGF3 complex translocates to the nucleus and binds to a consensus sequence known as interferon stimulated regulatory element (ISRE). Binding ISREs promote the of expression of many ISGs to limit viral infection [50]. However, herpesviruses encode multiple mechanisms to manipulate the innate immune response to not only prevent clearance. HCMV combats elimination by the innate immune response by terminating the signaling cascade at multiple checkpoints during a replicative infection [51-53]. Infection of fibroblasts and epithelial cells with the high-passage HCMV strain, Towne, causes proteasomal degradation of JAK1 to terminate phosphorylation of STAT1 [52, 54]. In addition, IE1 has been found to interact with STAT1 and STAT2 and prevent DNA binding of the ISGF3 complex to prevent ISG54 and Mx1 expression [55]. However, these studies were performed with a laboratory-adapted strain of HCMV that lacks genes in the UL*b’* locus, including UL138. Beyond shutting down parts of the signaling cascade, HCMV has been reported to stimulate pY701-STAT1 [56] and hijack the innate immune DNA sensor, IFI16, to activate the viral Major Immediate Early Promoter (MIEP) [57]. In addition, exogenous IFN*β*addition has been shown to reversibly silence the MCMV genome [48]. Further, HCMV latently-infected CD14+ monocytes exhibit a pro-inflammatory profile of gene expression, but are blunted in their response to type I or II triggers of STAT1 signaling [58]. Therefore, despite the anti-viral, nature of a type-I IFN response, HCMV likely commandeers this response for persistence [59].

Here we have demonstrated a mechanism by which UL138-host interactions stimulate pSTAT1 for the establishment of latency. Notably, we have found that UL138 interacts with UAF1 and USP1 (Table 1, Fig. 1). The scaffold protein UAF1 forms a complex with USP1, USP12, and USP46 that is essential for their activity. UAF1-USP1 in best understood for its role in regulating nucleation of DNA damage responses [29, 30, 36]. The UAF1 (WDR48) interaction with UL138 has been previously reported [34, 60]. While Nobre et al. [60] did not follow up this interaction, Li et al. [34] reported that UAF1 (WDR48) was important for viral genome synthesis and virus replication in MRC5 cells infected with the Han strain. Contrary to this finding, we find that inhibition of USP1 with C527 increased TB40/E genome replication in CD34+ HPCs, suggesting that at least the UAF1-USP1 complex is suppressive to viral genome replication, consistent with the role of UL138 in suppressing replication [17]. While we did not directly test the role of the UAF1 scaffold in HCMV infection, its knockdown is likely to have pleiotropic effects as it would result in the combined loss of USP1, USP12, and USP46 activity. In HPV infection, viral helicase, E1, promotes UAF1-USP1 activity for viral genome replication [61, 62]. The high-risk HPV oncogenes, E6 and E7, delay UAF1-USP1 mediated deubiquitination of the Fanconi anemia (FA) DNA crosslink repair, contributing to genomic instability [63]. Further studies are required to address the mechanisms by which UL138-UAF1-USP1 regulate HCMV replication and genome synthesis.

USP12 also co-precipitated with UL138 (Table 1), suggesting that in addition to USP1, that UL138 may also regulate USP12. The UAF1-USP12 complex has been implicated in sustaining pY701-STAT1 by binding to CREB-binding protein (CBP) and preventing acetylation of pY701-STAT1 and subsequent dephosphorylation by protein tyrosine phosphatase non-receptor type 1 (PTPN2) [64]. Although the role of the UL138-UAF1-USP12 complex will be addressed in future studies, the inclusion of USP12 in the IP/MS data set suggests an additional mechanism by which UL138 sustains pY701-STAT1. In support, JAK1 was also found a low-ranking hit identified in the IP/MS and bolsters the conclusions that UL138 interacts with host proteins to sustain STAT1 signaling.

UAF1-USP1 has been reported to sustain pY70-STAT1 signaling by deubiquitinating and preventing proteasomal degradation of TBK1 [40]. While we show that pY701-STAT1 depends on UL138 and USP1 at late times during infection in fibroblasts (Fig. 2-3), we were surprised that neither total or phosphorylated levels of TBK1 were affected by the absence of UL138 or USP1 as a possible mechanism to sustain pY701-STAT1 (Fig. 2C). This suggests that while both our studies and those of Yu et al. [40] demonstrate a role for USP1 sustaining pY701-STAT1, it may not be through the rescue of TBK1 from turnover. It has been reported that STAT1 must be deubiquitinated at Lys511 and Lys652 to license its recruitment to IFNAR and subsequence phosphorylation by JAK1/2 [65]. Therefore, it is possible that the UAF1-USP1 complex might deubiquitinate STAT1 to promote its interaction with and subsequent activation by IFNAR. It will be important to determine if this is the mechanism by which USP1 and UL138 function to sustain STAT1 signaling. Another study found that specifically pY701-STAT1 could be targeted for proteasomal degradation in the nucleus as an additional method to terminate pY701-STAT1 signaling [66]. This highlights another potential mechanism of UL138-USP1 in sustaining pY701-STAT1 during HCMV infection. Future studies will investigate these possibilities, as well as the interaction with USP12, to define the mechanisms by which UL138-UAF1 complexes sustain STAT1 activity during HCMV infection.

Contrary to our findings, Kalejta and colleagues showed that UL138 inhibits the innate immune response by targeting STING for degradation [41]. UL138 was shown to target STING, but this effect was only evident when both UL138 and STING were transiently overexpressed. Consistent with our findings (Fig. 2C), no UL138-dependent changes in STING protein levels were observed during infection. However, Kalejta and colleagues went on to show increased IFNβ and CXCL10 transcripts in *UL138*-M16_STOP_ infection in CD34+ HPCs, concluding that UL138 functions in immune evasion, degrading STING to abate a type-1 interferon response. However, an alternative explanation of the increase in IFNβ and CXCL10 expression is derived from the fact that transcripts were analyzed from unsorted hematopoietic cells. As UL138-mutant viruses are more replicative, increased levels of ISGs in a mixed population of infected and uninfected cells may be due to the enhanced replication of the mutant virus or the response of uninfected cells in the culture to secreted type 1 IFNs. Our results are consistent with that of the Kalejta laboratory in that changes in innate signaling appear to occur independently of changes in total or phosphorylated levels of TBK1 and STING.

We show that the prolonged pY701-STAT1 stimulated during HCMV infection correlates with an enhanced and sustained expression of several type 1 IFN-associated ISGs in a UL138-dependent manner. These include MX1, OAS, IFIT1, and IFIT2 (Fig 4B). Type 2 interferons also lead to the phosphorylation of STAT1, which form homodimers which bind to gamma activated sequences (GAS) to drive expression of ISGs that are similar and distinct to type 1 IFN driven ISGs [44]. However, we found no induction of a type-2 IFN driven ISG, ICAM1, above uninfected levels. Therefore, elevated pY701-STAT1 levels are most likely part of the type 1 IFN response. In both WT and Δ*UL138*_STOP_, the induced type 1 IFN ISGs diminish by 72 hpi despite sustained pY701-STAT1. This is undoubtedly due, in part, to robust inhibition of type-1 IFN signaling by multiple HCMV gene products [53, 67-70]. However, we also show that pY701-STAT1 localizes to nuclear viral RCs (Fig 5A, SI Fig. 2,) and binds to the viral genome to stimulate UL138 expression (Fig. 6). With respect to regulation of viral gene expression, STAT1 signaling been shown suppress lytic reactivation of MHV-68, similarly to UL138 but through a different mechanism. The MHV-68 ORF50 promoter, which is essential for the lytic switch, encode STAT1 repressive elements to negatively regulate lytic replication in response to IFNγ [71], whereas our findings suggest that HCMV commandeers STAT1 to stimulate the expression of a pro-latency gene—achieving the same end as MHV-68, but through a distinct mechanism. Further, it is possible that sequestration of pSTAT1 in viral RCs during HCMV infection, may also result in a distinct or diminished expression profile of ISGs at late times in infection.

UL138 has been shown to contribute to latency by stimulating the expression of the host transcription factor, EGR-1 [18], through MEK/ERK signaling. EGR-1 binds the viral genome upstream of UL138 to stimulate its expression for latency. Furthermore, inhibition of JAK1/2 activity in HCMV infected CD14+ monocytes increases viral transcripts [3]. Similarly, our findings here show that UL138 interaction with USP1 also regulates UL138 gene expression through sustaining activity of STAT1. Inhibition of EGFR or STAT1 with chemical inhibitors results in a loss of latency and increases virus replication in the absence of a stimulus for reactivation. Future studies will investigate more globally the changes in viral gene expression driven by EGR-1 and pY701-STAT1 to influence the latency program in CD34+ HPCs.

Viruses excel at manipulation of host signaling and innate pathways to evade elimination. However, as illustrated by our study, viruses also commandeer host defenses to their benefit, for example, in the case of viral persistence. Inhibition or loss of type-1 IFN signaling prevents the establishment of HIV-1 latency in macrophages [72] and murine gammaherpesvirus 68 (MHV-68) latency in splenocytes [73]. Epstein-Barr virus (EBV) latent membrane protein 1 (LMP-1) stimulates STAT1 activity mediated through NFкB signaling [74]. Further, EBV EBNA1, which is important for the maintenance of the EBV episome in latency, was shown to enhance the expression of STAT1 [75]. This induction sensitized cells to IFNγ treatment, although basal levels of ISGs were not changed, possibly due to LMP-2-suppression of STAT1 signaling. STAT1 has also been shown positively regulate the BamHI-Q promoter since interference with Jak/STAT signaling reduces BamH1-Q promoter activity [76]. In the context of EBV transformed B cells, STAT1 was shown to be important to the latency III viral program and to differentially regulate viral gene expression in tumors [77, 78]. Downregulation of STAT1 in this context resulted in increased viral lytic gene expression and lytic cycle entry, which is accompanied by downregulation of major histocompatibility complexes (MHC) class I and II at the cell surface. Interferon regulatory factors (IRF) also play complex roles in regulating viral gene expression. Interferon regulatory factor 2 (IRF2) and 7 (IRF7) bind and repress the BamHI-Q promoter of EBV to promote type III latency establishment [79, 80]. IRF7 is induced in a TRAF6- and RIP1-dependent manner by LMP-1 of EBV [81-84]. By contrast, IRF8, which is primarily expressed in hematopoietic cells, has been shown drive the expression of EBV lytic genes, such as BGLF2, and that the PIAS1 negative regulator of STAT1 signaling is important for latency [85]. In alphaherpesviruses, the viral E3 ubiquitin ligase, ICP0, of herpes simplex type 1 antagonizes STAT1 for replication. In the absence of ICP0, STAT1 restricts virus replication, suggesting a role for STAT1 in the establishment of HSV-1 latency [86]. STAT1 could regulate expression of the HSV-1 latency associated transcript (LAT), as binding sites have been mapped [87] and IFNβ modulates LAT expression and neuron survival [88]. While herpesviruses appear to largely commandeer innate signaling for the establishment of latency or to restrict lytic reactivation, it is also clear that this must be carefully controlled and some latency-associated viral factors may inhibit innate responses, such as MVH-68 M2 downregulating both STAT1 and STAT2 [89].

We have shown that the HCMV latency protein UL138 forms a unique interaction with UAF1-USP1 to control a typically antiviral response and utilize pY701-STAT1 for suppression of viral replication, at least in part, by stimulating of pro-latency gene expression programming. We have demonstrated the ability of UL138 to sustain pSTAT1 in fibroblasts and the importance of UL138-USP1 and STAT1 signaling in latency. It will be important to investigate the interplay between the UAF1-USP1 regulation of the DNA damage response and the innate immune responses in HCMV infection. While UL138 may modulate STAT1 activity to regulate innate responses for latency, it is also possible that STAT1 plays other critical roles central to latency and reactivation, such as its lesser understood role in hematopoietic differentiation that may occur independently of ISGs [90-94] or as a transcription factor recruited for viral gene expression. It will also be important to differentiate these possibilities in future work and to understand the role of other UAF1-USP complexes in the establishment of latency.

## Materials and Methods

### Cells

MRC-5 lung fibroblasts (ATCC), HEK293T/17 cells (ATCC), Sl/Sl stromal cells (Stem Cell Technology), M2-10B4 stromal cells (Stem Cell Technology), and CD34^+^ HPCs were maintained as previously described [19, 26]. Human CD34^+^ HPCs were isolated from de-identified medical waste following bone marrow isolations from healthy donors for clinical procedures at the Banner-University Medical Center at the University of Arizona. Latency assays were performed as previously described [19, 26]. In chemical treatments of MRC-5 lung fibroblasts, 0.88uM C527 (ApexBio), 5uM Ruxolitinib (STEMCELL Technologies) every 24 hours, or 1000U/mL universal type 1 interferon (R&D Systems) for 30 minutes. CD34+ HPCs were treated with 1uM C527 or every 24 hours with 500nM Ruxolitinib.

### Viruses

Bacterial artificial chromosome (BAC) stocks of TB40/E WT virus, a gift from Christian Sinzger [95], was engineered to express GFP from a SV40-promoter [17]. Creation of Δ*UL138*_STOP_ virus is as previously characterized [37].

### Plasmids and siRNAs

USP1_HA_ was purchased from addgene (BC050525). UL138_myc_ was overexpressed in pLVX_TetONE_puro vector and UAF1_HA_ was expressed from a pCIG vector. siGenome SMARTpool siRNAs targeting USP1 and nontargeting control (NTC) were purchased from Dharmacon. siRNAs were reverse transfected [96] into fibroblasts using Lipofectamine RNAiMAX reagent (Thermo Fisher Scientific) according to the manufacturer’s instructions. The next day, media was exchanged. 48 hours later, fibroblasts were infected (MOI=1) for 24, 48, and 72 hpi.

### Immunoblotting

Lysates were separated by electrophoresis on precast 4-20% precast gels (BioRad). Gels were transferred onto Immobilon-P PVDF membrane (EMD Millipore). Antibodies were incubated in with a blocking solution of 5% BSA in TBS-T, as per antibody manufacturer specifications. After antibody staining, blots were incubated with fluorescent secondary antibodies and imaged and quantitated using a Li-Cor Odyssey CLx infrared scanner with Image Studio software. Antibodies and sources are defined in Table 3.

### Immunofluorescence

Cells were grown on coverslips and infected at an MOI of 1 with TB40/E or Δ*UL138*_STOP_ virus. Uninfected cells were treated with DPBS or 1000U/ml of Universal Type 1 Interferon (R&D Systems) for 30 minutes. At each time point, cells were prepared for immunofluorescence by the cell signaling protocol for the pY701-STAT1 antibody. pY701-STAT1 and UL44 were detected through monoclonal antibodies and secondary antibodies were conjugated to Alexa Fluor 546 (green) or 647 (red). Coverslips were imaged using a DeltaVision deconvolution microscope.

### Subcellular fractionation

Fractionation was performed as described previously [97]. Fibroblasts were infected (MOI=1) and trypsinized and washed in PBS at each time point. Cells in suspension were then sequentially lysed in buffers containing detergents to separate subcellular compartments: 25 μg/mL digitonin (cytosolic fraction), 1% NP-40 (membrane-bound fraction), and 0.5% sodium deoxycholate and 0.1% SDS with 25 U/mL Benzonase® Nuclease (nuclear fraction).

### RT-qPCR and qPCR

Cells were infected with 1 MOI of TB40/E_GFP_ and DNA and RNA was isolated using Quick-DNA/RNA miniprep kit (Zymo Research) from 0–72 hpi. RNA was reverse transcribed into cDNA using SuperScript VILO cDNA Synthesis kit (Thermo Fisher Scientific). cDNA and DNA was quantified using LightCycler SYBR Mix kit (Roche) and corresponding primers (Table 2). Assays performed on Light Cycler 480 and corresponding software. Relative expression was determined using the ΔΔCT method normalized to H6PD. Viral genomes were quantified using a standard curve and normalized to RNAseP. For quantification of ISGs, fibroblasts were infected (MOI=1) for 24, 48, and 72 hpi. RNA was isolated using Quick-DNA/RNA miniprep kit (Zymo Research) and cDNA was prepared using 1000 ng of total RNA and random hexamer primers (Thermo Fischer Scientific). Real-time qPCR (TaqMan) was used to analyzed cDNA levels with specific TaqMan primer and probe sets (Thermo Fisher Scientifc). Relative expression was determined using the ΔΔCT method using GAPDH as the standard control.

**Table 2.**
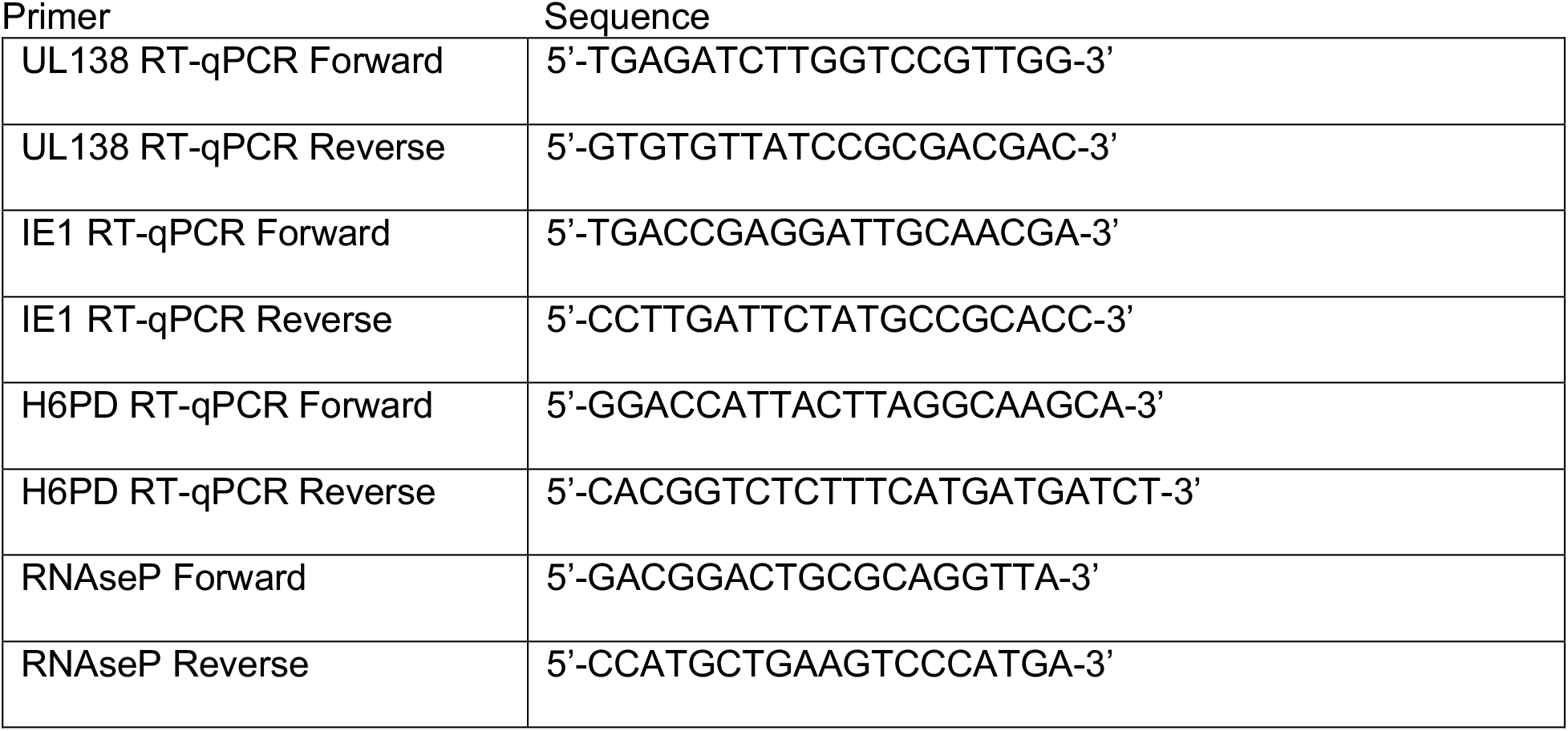
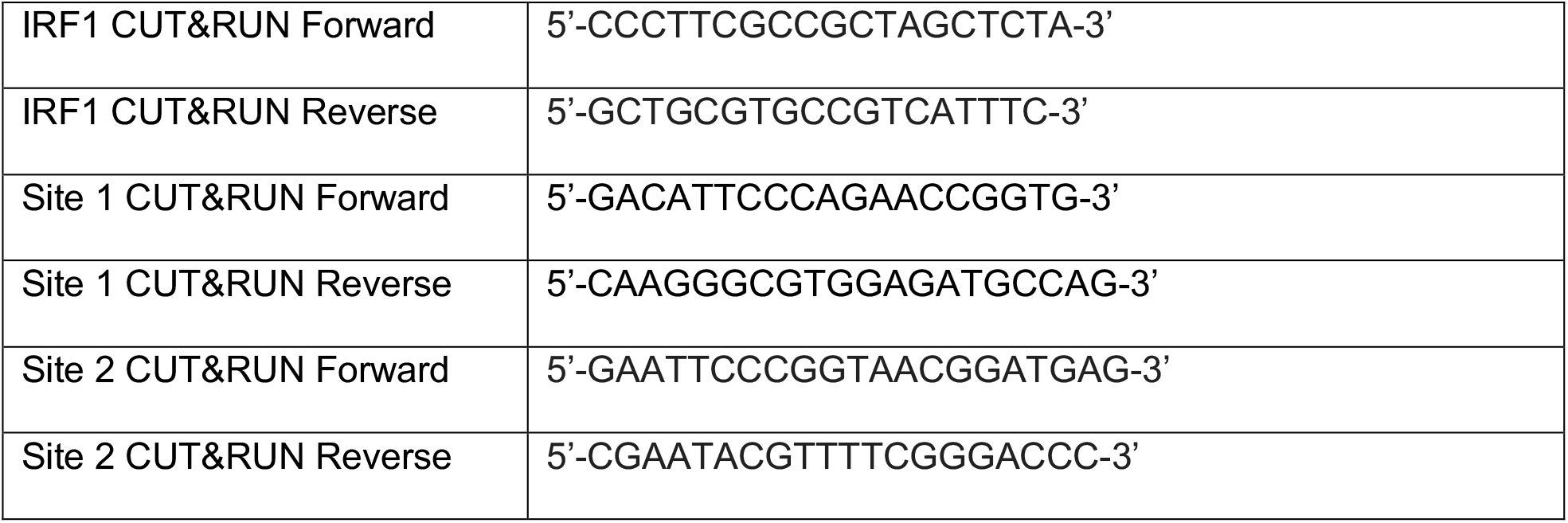
Primer sequences.

**Table 3.**
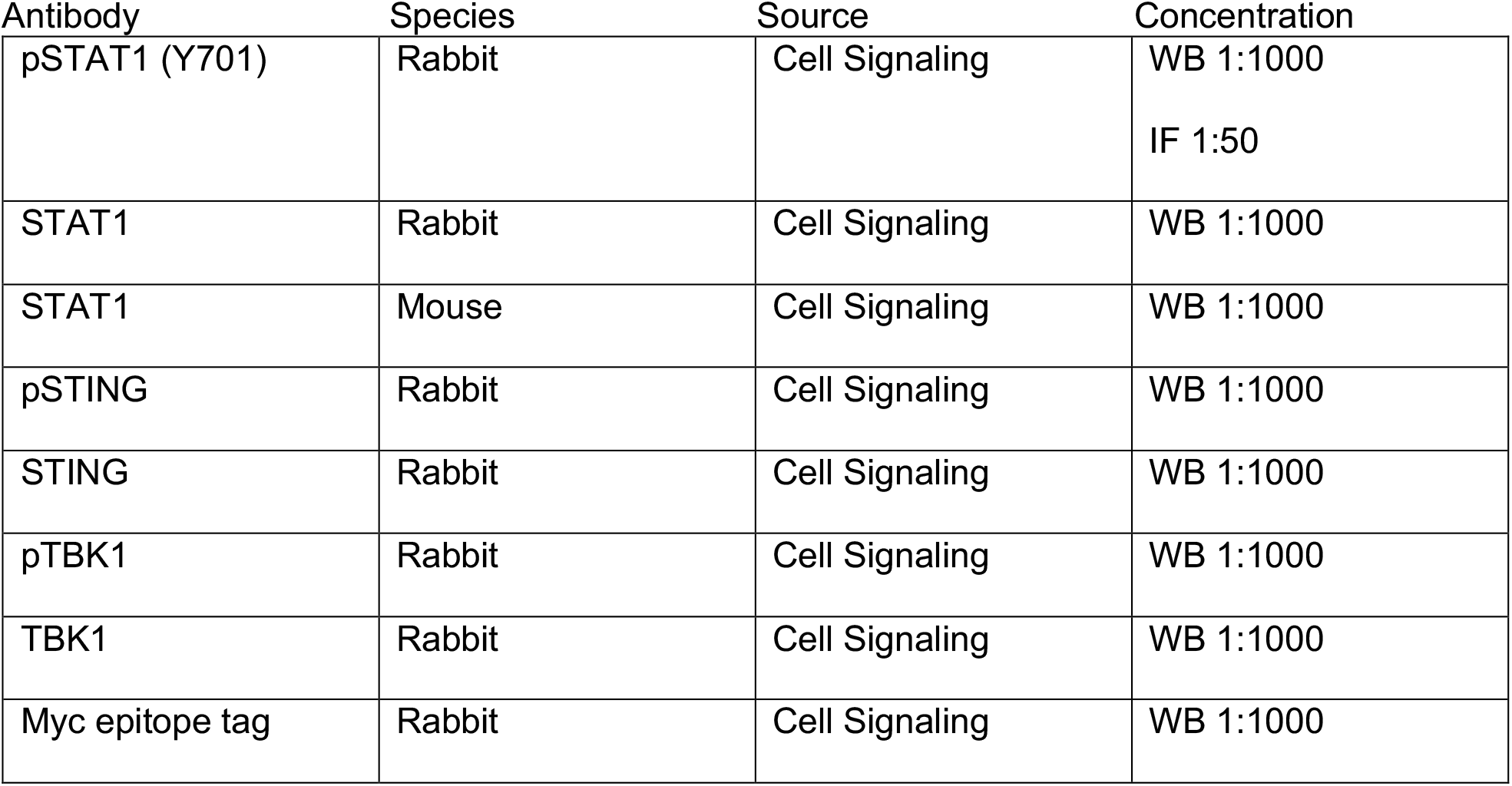
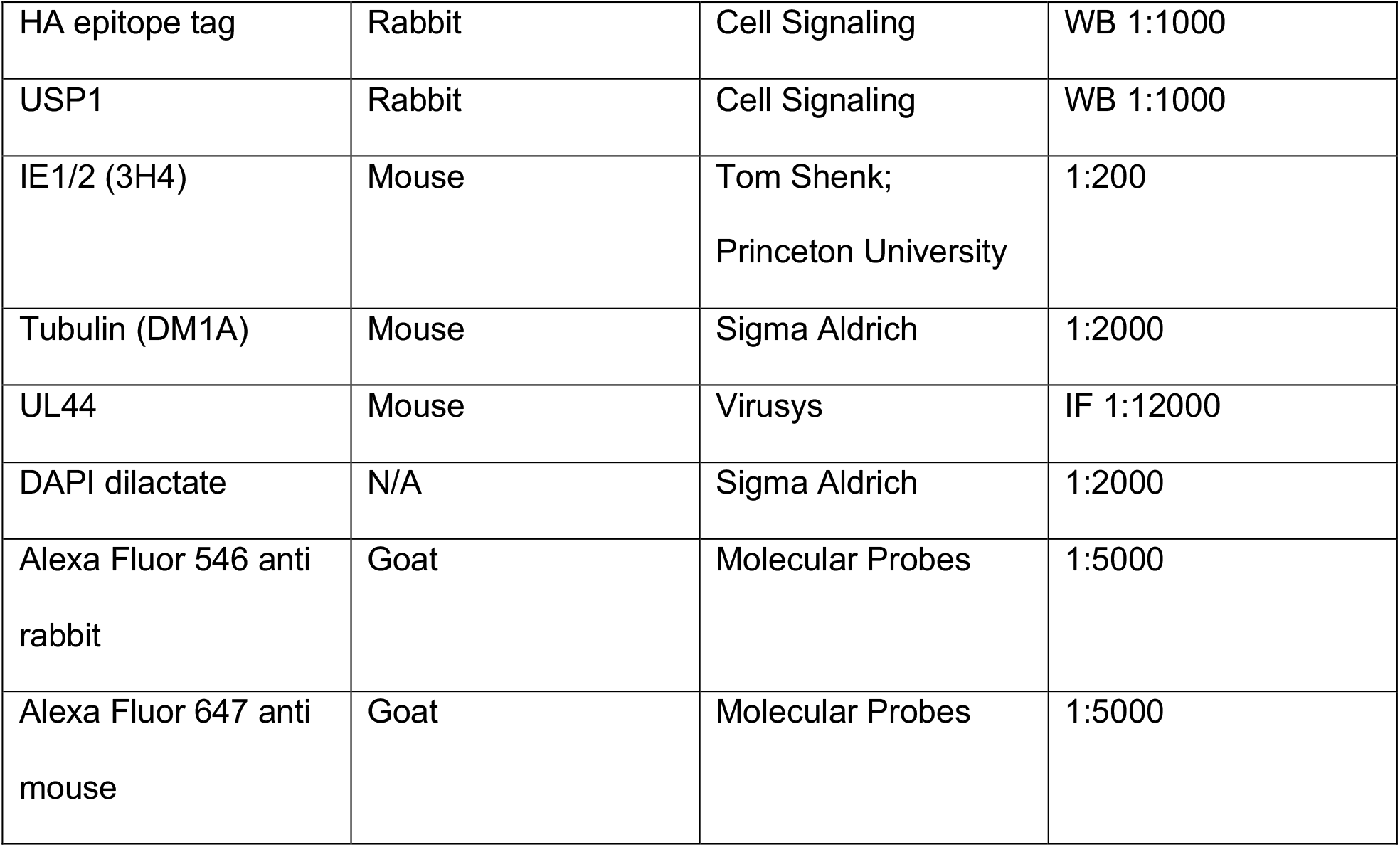
Antibody description and sources.

### CUT&RUN

For fibroblasts, 1e5 cells were utilized per condition. Cells were infected (MOI=1) with WT or UL138STOP virus for 48 hours and then processed for CUT&RUN (Cell Signaling Technologies) as per manufacturer’s recommended protocol. ChIP was carried out using pY701-STAT1, Histone H3 (positive control), and normal mouse (negative control). Isolation of DNA was performed through phenol chloroform extraction as per the manufacturer’s protocol. qPCR was performed with LightCycler SYBR Mix kit (Roche) and primers to pY701-STAT1 binding sites 1, 2, and IRF1. Relative expression was calculated against a 2% input control.Samples were then normalized to uninfected cells stimulated with universal type 1 interferon (R&D Systems).

### Viral Latency and Reactivation

Infectious centers were quantitated in CD34^+^ HPCs, as described previously (39). Frequency of infection centers were calculated using extreme limiting dilution analysis (42). For the investigation of USP1 activity during latency establishment, CD34^+^ HPCs were treated with 1uM/mL C527 (ApexBio) after sorting for CD34^+^ GFP^+^ populations and when stromals were replaced at 6 dpi. The study for USP1’s role in reactivation, 1uM/mL C527 was added at the time of reactivation. Proliferation of CD34^+^ cells during chemical inhibition was calculated by observing the fold change in the number of cells prior to and after inhibition for each condition. To understand JAK1/2’s role in latency establishment, CD34+ cells were treated with 500nM Ruxolitinib (STEMCELL Technologies) every 24 hours for 5 days. Cells were lysed and DNA was extracted at days 1 and 5 utilizing the Quick-DNA/RNA Miniprep (Zymo Research). Viral genomes were quantified using a standard curve and normalized to RNAseP.

### Statistical Analysis

All statistics were calculated using GraphPad Prism version 7 software. Statistics for experiments in this study were calculated using either unpaired student T-test or analysis of variance (ANOVA) for statistical comparison, which is indicated in the figure legends with asterisks representing statistical significance (* p-value <0.05, *** p-value <0.01, *** p-value <0.001).

### Data Availability

All data is contained within the manuscript. Proteomic data (peptide sequences) is supplied in Supplementary Information.

## Acknowledgements

We are grateful for the support of the Flow Cytometry and Human Immune Monitoring Shared Resource, supported through the Cancer Center Support Grant P30 CA023074 awarded to University of Arizona. We acknowledge Mark Curry and the Arizona Cancer Center/Arizona Research Laboratories Division of Biotechnology Cytometry Core Facility for expertise and assistance in flow cytometry. We acknowledge Dean Billheimer for statistics consultation. Special thanks to the Terry Fox Laboratory for providing the M2-10B4 and Sl/Sl cells. We acknowledge Dr. Tom Shenk for the gift of antibodies. This work was funded by grants from the National Institute of Allergy and Infectious Diseases to FG (AI079059, AI169728, and AI127335) and JAN (AI127335). KZ was funded by T32 (AG058503) from the National Institute of Aging.

PL is funded by AI079059-14S1.

